# The *C. elegans* gustatory receptor homolog LITE-1 is a chemoreceptor required for diacetyl avoidance

**DOI:** 10.1101/2025.04.20.649642

**Authors:** Alan Koh, Eduard Bokman, Alexey Gavrikov, Javier Rodriguez, Changchun Chen, Alon Zaslaver, André EX Brown

## Abstract

The nematode *C. elegans* does not have eyes but can respond to aversive UV and blue light stimulation and even distinguish colours. The gustatory receptor homolog LITE-1 was identified in forward genetic screens for worms that failed to respond to blue light stimulation. When LITE-1 is expressed in body-wall muscles, it causes contraction in response to blue light suggesting that LITE-1 is both necessary and sufficient for blue light response. Here we show that in addition to light avoidance, LITE-1 is also required for worms’ avoidance of high concentrations of diacetyl, an odorant that is attractive at low concentrations. Like blue light, diacetyl causes muscle contraction in transgenic worms engineered to express LITE-1 in body-wall muscles. These data are consistent with a direct chemoreceptor function for LITE-1 which would make it a multimodal sensor of aversive stimuli.

## Introduction

Sensory perception is fundamental to how organisms interact with their surroundings. Animals, including *Caenorhabditis elegans*, detect and respond to a wide range of environmental cues such as light and odorants. Despite having a compact nervous system of only 302 neurons, *C. elegans* exhibits sensory capabilities that allow it to effectively navigate and adapt to its environment (Bargmann, 2006; Cook et al., 2019). Odorants such as volatile organic compounds are detected via specialised olfactory receptor neurons. These neurons express specific receptors that bind to specific molecules and trigger distinct behavioural responses (Bargmann, 2006; Bargmann et al., 1993; Bargmann & Horvitz, 1991). Diacetyl, a volatile organic α-diketone found in *C. elegans*’ natural environment elicits opposing behaviours depending on its concentration. At low concentrations, diacetyl is sensed by the olfactory receptor ODR-10, promoting attraction (Itskovits et al., 2018; Sengupta et al., 1996; Zhang et al., 1997). In contrast, higher concentrations induce an avoidance response, thought to be mediated by the receptor SRI-14 (Taniguchi et al., 2014).

In addition to chemical cues, *C. elegans* also detects and responds to light, despite lacking eyes or classical photoreceptors. This light sensitivity is mediated by LITE-1, a member of the Gustatory Receptor family (Montell, 2009). LITE-1 was first identified through genetic screens for mutants defective in light avoidance behaviour (Edwards et al., 2008). Unlike canonical photoreceptors such as rhodopsins or opsins, LITE-1 shares little sequence similarity with known light sensing proteins (Edwards et al., 2008; Gong et al., 2016; Liu et al., 2010). Further underscoring its uncanonical nature, LITE-1 adopts an inverted membrane topology that is unlike classical G protein-coupled receptors (GPCRs) (Gong et al., 2016).

LITE-1 phototaxis behaviour is driven by multiple light responsive sensory neurons, including ASJ, ASH and ASK (Liu et al., 2010; Ward et al., 2008; Zhang et al., 2022). Signal transduction occurs via G protein signalling, upregulation of cGMP levels and activation of cyclic nucleotide-gated (CNG) channels, leading to a rapid avoidance response (Edwards et al., 2008; Liu et al., 2010; Ward et al., 2008). Recent studies suggest that LITE-1 may function as a light activated ion channel with a putative ligand binding pocket that is activated by ultraviolet and blue light via key tryptophan residues (Edwards et al., 2008; Gong et al., 2016; Hanson, Scholüke, et al., 2023; Ward et al., 2008).

Here we show that beyond its role in photoreception, LITE-1 is also required for the avoidance of high concentrations of diacetyl and that diacetyl induces muscle contraction in transgenic worms expressing LITE-1 in body-wall muscles. Our findings support the notion that LITE-1 functions as a multifunctional receptor involved in both light and chemical sensing.

## Results

### LITE-1 is required for diacetyl and blue light avoidance

A recent screen of a panel of worm strains that collectively contained knockouts of all non-essential GPCRs identified a triple mutant that failed to avoid high concentrations of diacetyl (Pu et al., 2023). One of the mutated genes was *lite-1*. We performed a chemotaxis assay to test single mutants of several loss-of-function alleles of *lite-1* and found they all failed to avoid high concentrations of diacetyl (Fig. 1A and 1B) while remaining attracted to lower concentrations (Fig. 1C). As LITE-1 is sensitive to light, we next examined whether ambient light affected these responses by comparing chemotaxis under light and dark conditions. Under both conditions, wild-type animals strongly avoided high concentration of diacetyl, whereas *lite-1* mutants still failed to do so, indicating that this phenotype is light independent (Fig. S1A). As expected, these *lite-1* mutants also no longer respond to blue light stimulation (Fig. 1D) (Edwards et al., 2008).

**Figure 1.**
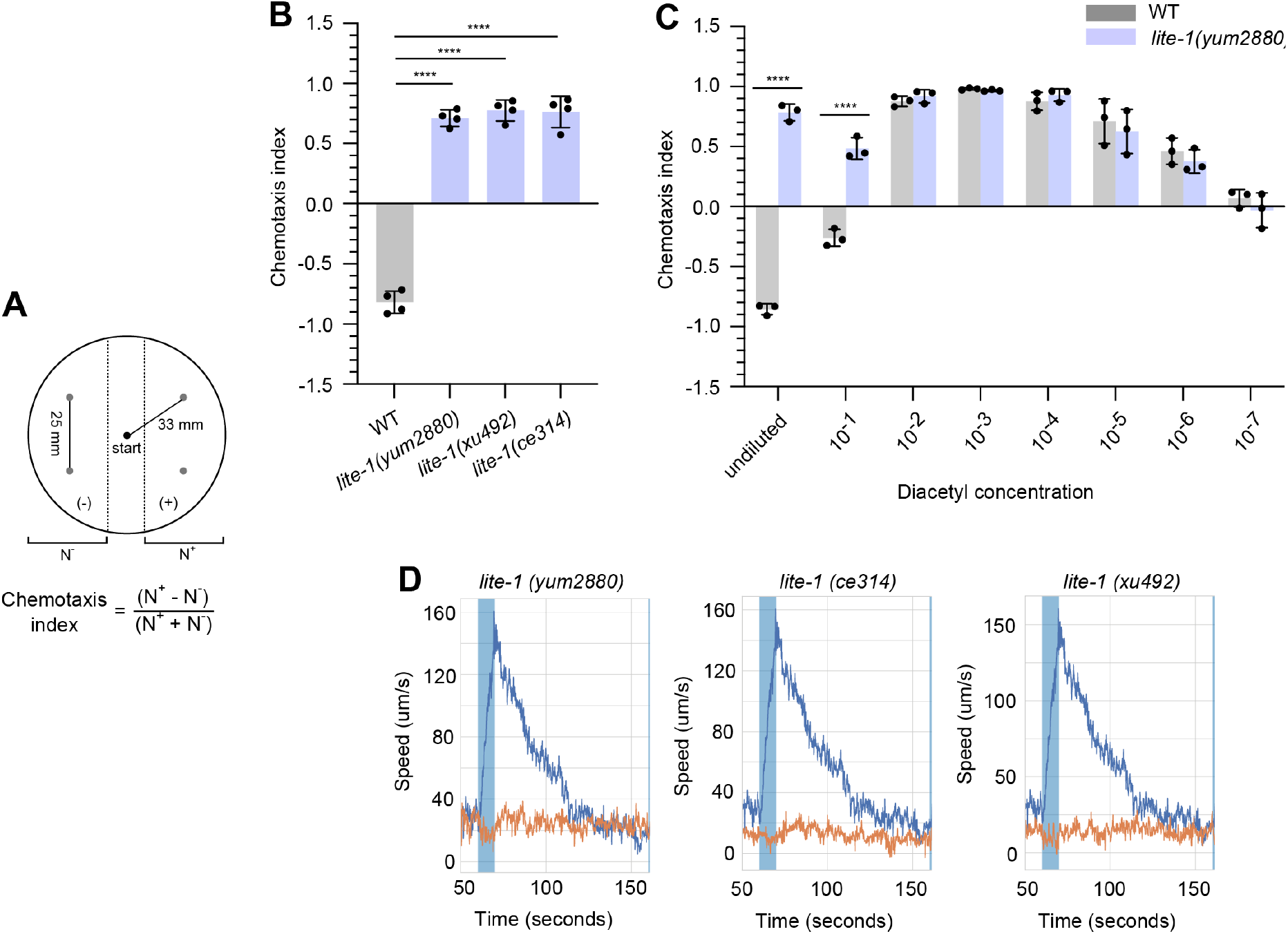
(A) Schematic of chemotaxis assay setup. (B) Chemotaxis index of wild-type animals and mutants with different alleles of *lite-1* that have defects in diacetyl response. P-values are from unpaired two-tailed t-tests, n = 4 biological replicates, each an average of 4 plates. (C) Chemotaxis index of wild-type and *lite-1 (yum2880)* animals with different diacetyl concentrations (5 µl). P-values are from unpaired two-tailed t-tests, n = 3 biological replicates, each an average of 4 plates. (D) Time series of wild-type (blue line) animals and mutants with different alleles of *lite-1* (orange line) when stimulated with blue light (blue shaded box). n = 3 biological replicates, each an average of 12 wells.

We next examined whether the *lite-1* mutants’ attraction to high concentrations of diacetyl depends on the low-concentration diacetyl receptor ODR-10 by analysing *lite-1; odr-10* double mutants. The double mutants remained attracted to high concentrations of diacetyl (Fig. S1B). This attraction may result from the activation of other, less specific odorant receptors whose signals become unmasked in the absence of LITE-1 mediated inhibition. This is consistent with earlier reports where worms do not completely lose attraction to low concentrations of diacetyl in the absence of ODR-10, suggesting the presence of additional low concentration receptors (Sengupta et al., 1996; Taniguchi et al., 2014). We also examined whether the avoidance behaviour depends on TAX-4 mediated chemosensory signalling. Like wild-type worms, *tax-4* mutants retained strong avoidance to high concentration of diacetyl, suggesting that TAX-4 independent pathways contribute to the avoidance response (Fig. S2). Consistent with additional aversive pathways, observations of *lite-1* mutants show that they do not reach the highest concentration of diacetyl (Supplementary video. 1 and 2).

### Chemosensory neurons ADL and ASK are involved in avoidance of diacetyl

To identify neurons involved in the response to high concentrations of diacetyl, we used calcium imaging to record 11 pairs of sensory neurons in worms exposed to a pulse of diacetyl (Fig. 2A). In *lite-1* mutants, the LITE-1-expressing sensory neurons ADL and ASK showed a decrease in calcium levels (Fig. 2B and S3A). Since ADL and ASK sense both attractive and aversive stimuli, it is plausible that the attractive response to diacetyl suppresses their activity. In the absence of LITE-1-mediated activation, the observed decrease in calcium responses may reflect the unopposed inhibitory signal input. In contrast, another LITE-1 expressing neuron, ASH, showed a normal response compared to wild-type animals, which is likely due to its expression of SRI-14 (Fig. 2B and S3A). SRI-14 has previously been shown to respond weakly to high concentration of diacetyl (Taniguchi et al., 2014). The response of the other tested neurons was not significantly different between *lite-1* mutants and wild-type animals (Fig. 2B and S3A). Additionally, we observed that the *unc-13* mutants, which are impaired in synaptic activity, had little effect on ADL response, but had a slower decay of the calcium signal in ASK neurons compared to wild-type animals (Fig. 2C and S3B) (Richmond et al., 1999). This supports the presence of additional inhibitory signals influencing ASK activity. Together, these data support the involvement of these neurons, although neural signalling through gap junctions or via neuromodulation by other receptors could not be ruled out.

**Figure 2.**
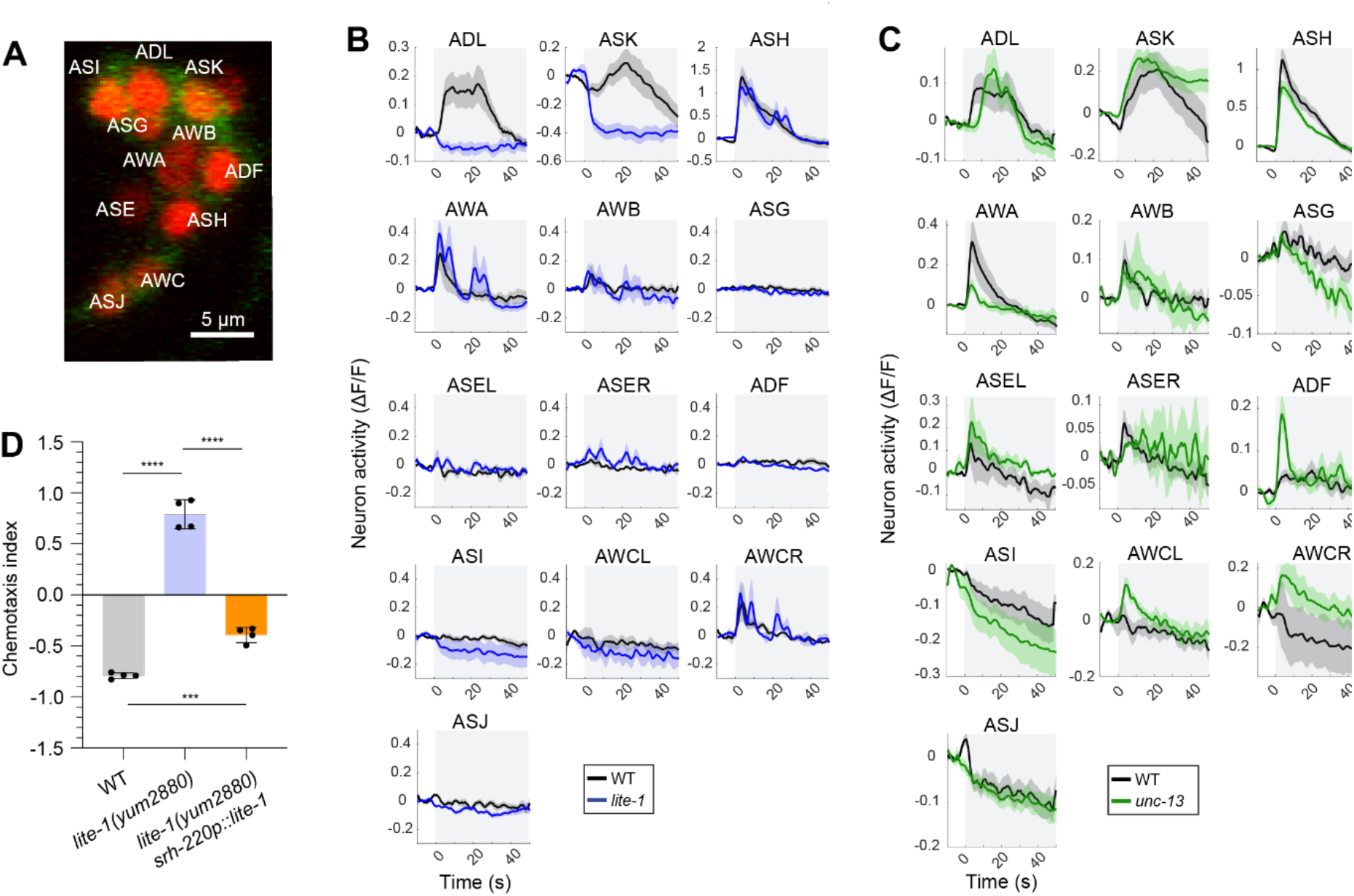
(A) Confocal image of labelled sensory neurons and calcium imaging traces showing the responses of three neurons when exposed to high concentrations of diacetyl. (B) Neuronal activity of wild-type controls with *lite-1* mutants. (C) Neuronal activity of wild-type controls to *unc-13* mutants. Grey shaded box denotes the presence of diacetyl. Shaded area around the activity dynamics denotes standard error. (D) Chemotaxis index of wild-type animals and rescue mutants expressing *lite-1* cDNA from the *srh-220* promoter in ADL neuron. P-values are from unpaired two-tailed t-tests, n = 4 biological replicates, each an average of 4 plates.

We next expressed *lite-1* genomic DNA under the ADL specific promoter *srh-220*, which restored the avoidance phenotype, although it is not a complete rescue of wild-type behaviour (Fig. 2D) (Hao et al., 2010). Together with calcium imaging data, this suggests that proper avoidance likely requires input from both ADL and ASK neurons.

### LITE-1 is sufficient for diacetyl response in other cell types

To test whether LITE-1 is sufficient for a response to high diacetyl concentrations, we used a transgenic strain that expresses LITE-1 in body-wall muscles (Edwards et al., 2008). Worms ectopically expressing LITE-1 in body muscle cells evoke muscle contraction and shortening of body length in response to blue light stimulation (Edwards et al., 2008). When these worms are exposed to a 1:10 dilution of diacetyl, they showed a rapid and near complete paralysis whereas wild-type animals continue to swim (Fig. 3A and 3B). These paralysed worms also showed significant contraction in average body length, contracting by ∼14.8% after diacetyl exposure (Fig. 3C). Consistent with muscle contraction, these worms also ejected eggs, while wild-type worms do not (Fig. 3D). To rule out contributions of neuronal LITE-1, the assay was repeated in *lite-1 (ce314)* mutants. In these strains, diacetyl still induced paralysis, muscle contraction, and egg ejection (Fig. S4). These results suggest that LITE-1 is activated by diacetyl. Although it remains possible that it is a diacetyl metabolite that activates LITE-1, this conversion would need to happen within seconds because maximal contraction occurs within 10 seconds of addition.

**Figure 3.**
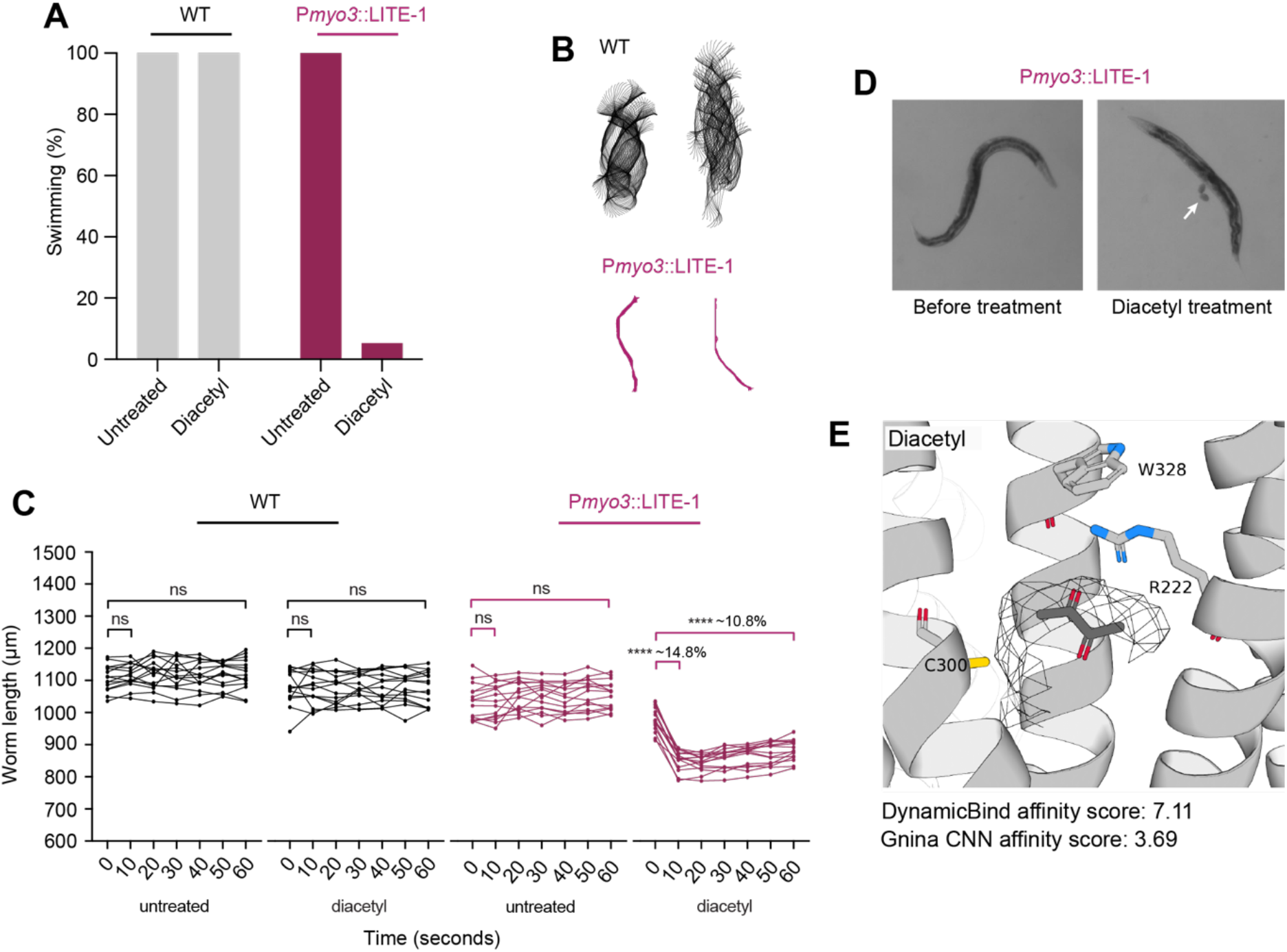
(A) Percent of worms that are swimming at ten seconds after exposure to buffer untreated or to diacetyl. Wild-type animals continue swimming in buffer and diacetyl (WT, left) whereas worms expressing LITE-1 in body wall muscles (*Pmyo-3::*LITE-1, right) are mostly paralysed when treated with diacetyl. (B) Sample midline trajectories from wild-type and *Pmyo-3::*LITE-1 animals showing difference in motion (each trajectory shows 6 seconds of swimming). (C) Worm body length over time after treatment. Wild-type animals do not show a significant length difference in either buffer or diacetyl treatments (left) whereas worms expressing LITE-1 in body-wall muscles show a significant contraction in response to diacetyl (insert showing the average contraction). P-values from paired two-tailed t-test, n ≥ 15 worms. (D) Image of a *Pmyo-3::*LITE-1 animal that ejected eggs within ten seconds of diacetyl treatment (white arrow). (E) Molecular docking of diacetyl into the AlphaFold-predicted structure of LITE-1 with predicted micromolar affinity.

### 2,3-pentanedione also interacts with LITE-1

We next tested whether *lite-1* is required for avoidance of other high concentration odorants. Of the seven odorants tested, *lite-1* mutants showed a defective response to 2,3-pentanedione, a chemical structurally related to diacetyl (Fig. 4A). The effect was weaker than that observed for diacetyl, with worms still avoiding high concentrations of 2,3-pentanedione but more weakly than wild-type animals, independent of ODR-10 (Fig. S1B). Supporting this observation and consistent with a ligand receptor interaction, *in silico* molecular docking predicts that both diacetyl and 2,3-pentanedione occupy the putative chromophore binding site, with diacetyl exhibiting the strongest affinity (Fig. 3E, 4B and S5) (Hanson, Scholüke, et al., 2023). Like diacetyl, 2,3-pentanedione also evokes paralysis, muscle contraction and egg ejection of worms expressing LITE-1 in body-wall muscle cells, independent of neuronal LITE-1 (Fig. 4C, 4D, 4E and S6).

**Figure 4.**
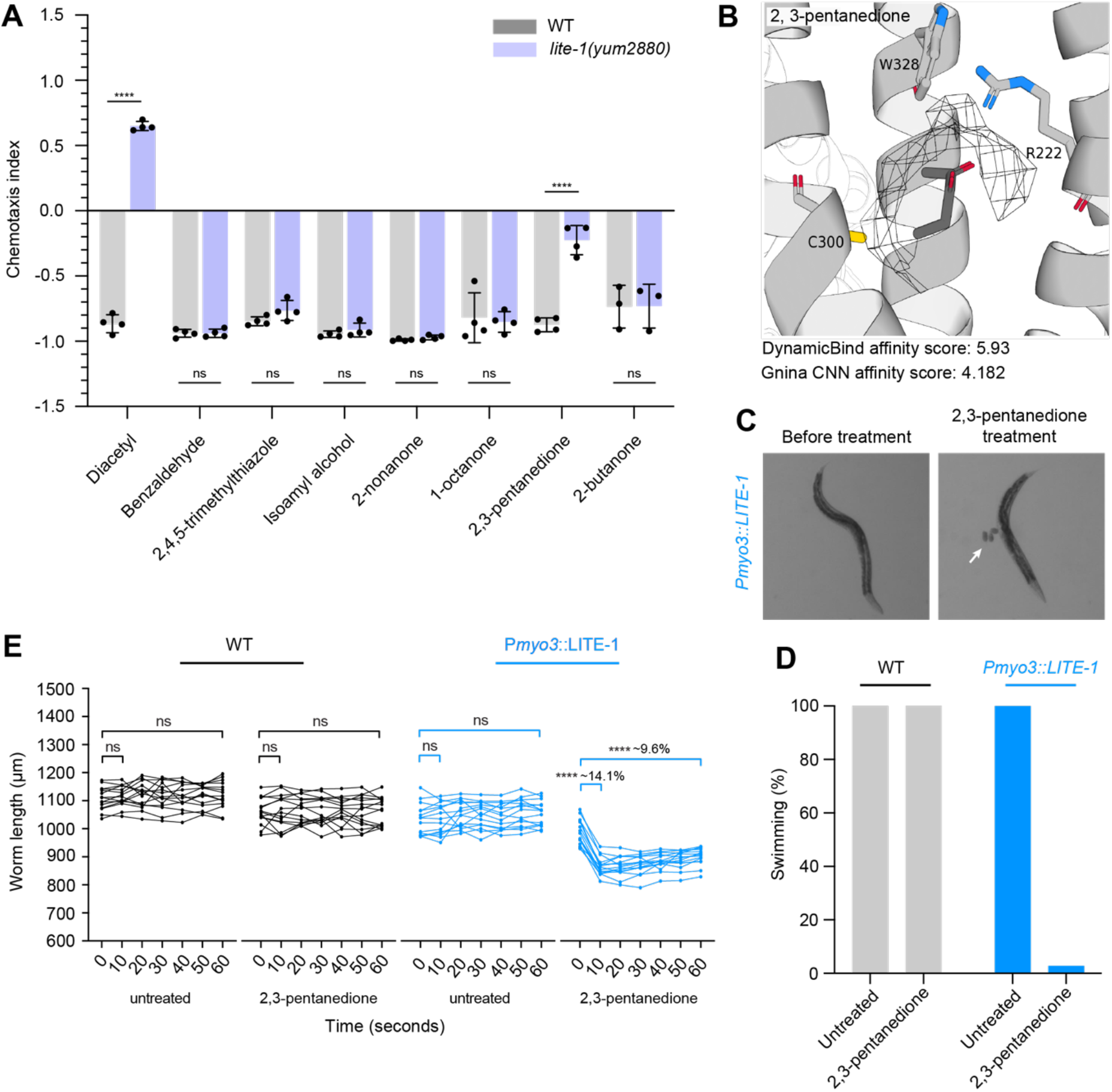
(A) Chemotaxis index of wild-type animals and *lite-1* mutants for known odorants. P-values are from unpaired two-tailed t-tests, n = 4 biological replicates each an average of 4 plates. (B) Molecular docking of 2,3-pentanedione into the AlphaFold-predicted structure of LITE-1 with predicted micromolar affinity. (C) Image of a *Pmyo-3::*LITE-1 animal that ejected eggs within ten seconds of 2,3-pentanedione treatment (white arrow). (D) Percent of worms that are swimming at ten seconds after exposure to buffer untreated or to 2,3-pentanedione. Wild-type animals continue swimming in buffer and 2,3-pentanedione (WT, left) whereas worms expressing LITE-1 in body-wall muscles (*Pmyo-3::*LITE-1, right) are mostly paralysed when treated with 2,3-pentanedione. (E) Worm body length over time after treatment. Wild-type animals do not show a significant length difference in either buffer or 2,3-pentanedione treatments (left) whereas worms expressing LITE-1 in body-wall muscles show a significant contraction in response to 2,3-pentanedione (insert showing the average contraction). P-values from paired two-tailed t-test, n ≥ 15 worms.

## Discussion

LITE-1 is best known for its role in light sensation in *C. elegans*, but how it senses light is not yet clear. In particular, an indirect mechanism in which light produces photoproducts that activate the channel has not been ruled out and *a priori* has always seemed plausible given that LITE-1 is similar to gustatory receptors that are known to sense chemicals (Montell, 2009). A direct diacetyl response would make a photoproduct-based mechanism more plausible, particularly as diacetyl has been demonstrated to act as a photosensitizer (Huang et al., 2019; Lian et al., 2022). What this photoproduct might be remains elusive, although reactive oxygen species could be involved. Indeed, light exposure can generate hydrogen peroxide in neurons, which has been proposed to inactivate activated LITE-1 (Bischer et al., 2023; Gong et al., 2016; Quintin et al., 2022). The identification of a second structurally related compound, 2,3-pentanedione may help in narrowing down photoproducts with similar moieties. Direct and photoproduct light sensing are not mutually exclusive. Based on LITE-1’s predicted structure and its absorption spectrum, it has recently been proposed that UV may be directly sensed while the blue light response may be mediated through a photoproduct or an unidentified chromophore (Hanson et al., 2023). While it is possible that UV absorption within cells may lead to diacetyl production, what this precursor might be remains elusive. Diacetyl generated in this way would need to be produced at a sufficient concentration to activate LITE-1. Based on the behavioural and muscle-contraction data, this occurs at relatively high concentrations.

While our study primarily focused on chemosensory neurons, it remains possible that non-chemosensory neurons expressing LITE-1 may contribute to diacetyl detection or modulating the behavioural response. Indeed, other LITE-1 expressing non-chemosensory neurons, such as the interneuron AVG, which mediates light sensing through the release of the neuropeptide NLP-10, as well as the phasmid neuron PHA and the pharyngeal interneuron I2, which are required for hydrogen peroxide sensing, are possible candidates. (Bhatla & Horvitz, 2015; Dunkel et al., 2025; Quintin et al., 2022). Nevertheless, phototransduction and chemotransduction appear to involve shared sensory neurons, such as ASK and ASH, suggesting a potential overlap in their underlaying signalling mechanisms (Zhang et al., 2022). It is possible that LITE-1 chemosensory response may rely on components necessary for LITE-1 photosensory function, such as G proteins (GOA-1, GPA-3), guanylate cyclase (DAF-11, ODR-1), phosphodiesterase (PDE-1,2,5) and neuropeptide signalling (NLP-10/NPR-35) (Dunkel et al., 2025; Liu et al., 2010; Ward et al., 2008). Additionally, while our molecular docking analysis supports the possibility of diacetyl and 2,3-pentanedione occupying LITE-1’s putative chromophore binding site, this remains to be experimentally validated. It would therefore be informative to test *lite-1* point mutants whose residues are predicted to be involved in ligand interactions, binding pocket properties, or affecting specific LITE-1 functions, such as light sensing or channel gating (Bischer et al., 2023; Hanson, Scholüke, et al., 2023).

Nevertheless, we have shown that LITE-1 is required for the avoidance of high concentrations of diacetyl and a related compound 2,3-pentanedione. Furthermore, we have shown that LITE-1 is sufficient to make body-wall muscle cells responsive to diacetyl and 2,3-pentanedione, consistent with a direct response. Diacetyl is an ecologically relevant molecule present in *C. elegans*’ natural environment that has been hypothesised to be involved in prey-finding since low concentrations produced by lactic acid bacteria on rotting fruit is attractive (Choi et al., 2016). Avoidance of high concentrations of diacetyl may therefore also be ecologically relevant. In this case, LITE-1 has likely evolved to be a multimodal receptor of aversive stimuli akin to other multimodal receptors such as TRPV1, which responds to heat, capsaicin, and acidic pH (Zhang et al., 2023).

## Material and methods

### Bacterial and worm strains maintenance

The *E. coil* wild-type isolate OP50 was sourced from Caenorhabditis Genetics Center (CGC) and maintained on lysogeny broth (LB) agar under standard laboratory conditions. The following worm strains were used in this study, N2; wild-type isolate (CGC), PR678; *tax-4(p678)*, CHS1173; *odr-10(yum2055)* (Pu et al., 2023), CHS1610; *lite-1(yum2880)* (Pu et al., 2023), CHS1721 *lite-1(yum2880) odr-10(yum2055)* (Pu et al., 2023), KG1180; *lite-1(ce314)* (Edwards et al., 2008), KG1271; *lite-1(ce314) cels37[myo-3*_*prom*_*::lite-1::GFP]* (Edwards et al., 2008), KG1272; *cels37[myo-3*_*prom*_*::lite-1::GFP]* (Edwards et al., 2008), TQ8245; *lite-1(xu492)* (Zhang et al., 2020), ZAS280; *In[osm-6::GCaMP3, osm-6::ceNLS-mCherry-2xSV40NLS]* (Iwanir et al., 2019), ZAS371; *unc-13(s69) In[osm-6::GCaMP3, osm-6::ceNLS-mCherry-2xSV40NLS]* (Bokman, Kalij, et al., 2024; Bokman, Pritz, et al., 2024), ZAS526; *lite-1(ce314) In[osm-6::GCaMP3, osm-6::ceNLS-mCherry-2xSV40NLS]*, were cultured on Nematode Growth Medium seeded with *E. coli* (OP50) as previously described (Brenner, 1974). Worm synchronization was performed by bleaching unsynchronised gravid adults (Barlow, 2019).

### Chemotaxis assay

The chemotaxis assays were conducted under ambient light, but care was taken to prevent exposure to any strong light source at 20°C in 9 cm plates filled with 10 ml of agar (1.7% agar, 1 mM CaCl_2_, 1 mM MgSO_4_ and 25 mM K_2_HPO_4_ pH 6). Two parallel lines, 1 cm apart, were drawn on the middle of the plates which define the area to place the animals. Two odorant spots were designated on one side of the plate, while two control spots were positioned on the opposite side (Fig. 1A). 1 μl of 1 M NaN_3_ was added to all four spots to immobilise the animals. Different volumes of each odorant: diacetyl (5 μl), 2,4,5-trimethylthiazole (7.5 μl), benzaldehyde (2.5 μl), isoamyl alcohol (7.5 μl), 2,3-pentanedione (10 μl), 2-nonanone (1 μl), 1-octanol (2.5 μl), 2-butanone (10 μl) and equal volume of ethanol (control) were spotted onto the respective dots. Synchronized day-one adults were washed 3 times in chemotaxis buffer (1 mM CaCl_2_, 1 mM MgSO_4_ and 25 mM K_2_HPO_4_ pH 6) before being transferred onto the centre of the assay plate where the assay will last for 1 hr at 20 °C. The chemotaxis plates were then stored at 4 °C before counting. The chemotaxis index was calculated as (number of animals on the odorant side – number of animals on the ethanol side) / (sum of all animals on both sides). For chemotaxis conducted in the dark, animals were maintained in darkness from hatching to adulthood. Brief exposure to dim ambient light occurred only during washing and transfer to agar plates, as the latter step had to be performed in a fume hood due to the odorant.

Following transfer, plates containing the animals were immediately covered with foil to maintain darkness.

### Paralysis assay

The paralysis assays were conducted in the dark (animals were kept in the dark from hatching to adulthood, with ambient light exposure minimised) at 20°C in 9 cm plates filled with 10 ml of agar (1.7% agar, 1 mM CaCl_2_, 1 mM MgSO_4_ and 25 mM K_2_HPO_4_ pH 6). Synchronized day-one adults were washed 3 times in chemotaxis buffer (1 mM CaCl_2_, 1 mM MgSO_4_ and 25 mM K_2_HPO_4_ pH 6) before being transferred onto the centre of the plate. Animals were allowed to crawl on the plate for five minutes before a thin layer of ∼;20 μl chemotaxis buffer containing either diacetyl or 2,3-pentanedione (1:10) was spread across the worms to completely cover them. Video acquisition began prior to buffer addition, recorded at 8 fps using a Basler acA4024-29um monochrome camera under red-filtered illumination, controlled with pylon Viewer 64-Bit (version 6.2.0.21487). Measurement of animal length was quantified manually using Fiji (Schindelin et al., 2012). Only worms that were in focus were included in the analysis.

### Blue light stimulation assay

The blue light stimulation assays were conducted at 20°C in 24-well plates containing agar (1.7% agar, 1 mM CaCl_2_, 1 mM MgSO_4_ and 25 mM K_2_HPO_4_ pH 6). Synchronized day-one adults were transferred onto the centre of the 24-well plate pre-seeded with OP50. The plates were then transferred onto the multi-camera tracker for 30 min to allow acclimatization to the tracker room in the dark before imaging. Videos were acquired at 25 frames per second using a shutter speed of 25 ms and a resolution of 12.4 μm/pixel. Videos were taken in the following order: 300 s pre-stimulus, blue light stimulus with one 10 s pulse of blue light at 60 s and a 100 s post-stimulus recording. Videos were then processed with Tierpsy Tracker as previously described (Barlow et al., 2022; Javer et al., 2018).

### Calcium imaging

Young adult worms were inserted into a microfluidic chip and anesthetized with 10 mM levamisole and allowed to habituate for 10 mins as previously described (Chronis et al., 2007). Recordings began with 1 min of light habituation, after which the stimulus was presented for 1 min The stimulus used was 1:5 diacetyl in CTX buffer (5 mM KH_2_PO_4_/K_2_HPO_4_ pH 6, 1 mM CaCl_2_ and 1 mM MgSO_4_). Recordings were made using a Nikon A1R + confocal laser scanning microscope with a water immersion × 40 (1.15NA) objective controlled by the Nikon NIS-elements software. Z-axis resolution was 0.5-0.8 μm. Acquisition rate was ∼;1.5 volumes/second, interpolated to a final framerate of 2 Hz.

### Neuron identification and signal extraction

Neuron segmentation and tracking was done on the nuclear mCherry signal using an algorithm developed by (Toyoshima et al., 2016). Neurons were identified visually based on anatomical positions. Neurons that could not be unambiguously identified were removed from analysis. Calcium readings were taken from within 0.9 of the radius of the segmented nuclear sphere. Signal intensity was normalized by baseline activity, defined as the lowest point of a 10-frame running average. All data analyses were performed using in-house MATLAB scripts (Pritz et al., 2023).

### Molecular docking

Molecular docking was performed in two stages. Diacetyl was first docked to the tetrameric LITE-1 model using DynamicBind without a predefined binding pocket (Hanson, Scholuke, et al., 2023; Lu et al., 2024). The generated complexes were then ranked using DynamicBind confidence score, and the highest-ranked poses were used to identify the candidate binding site. The top DynamicBind pose was then used to define the box region for redocking with Gnina (McNutt et al., 2021). Gnina poses were ranked using the CNN score. The PDB file containing a representative DynamicBind derived docking pose of diacetyl within the binding groove of LITE-1 is available at https://doi.org/10.6084/m9.figshare.32745054.

## Acknowledgements

We thank Na Yu and John Bergqvist for technical assistance and Denise Walker, Bill Schafer, and Alexander Gottschalk for providing strains and Marty Chalfie for useful discussions. This work was funded by the Medical Research Council (grant MC-A658-5TY30 to AEXB), the Israel Science Foundation (grant 1939/23 to AZ) and the Russian Science Foundation (grant 25-76-30006 to AG). Some strains were provided by the CGC, which is funded by NIH Office of Research Infrastructure Programs (P40 OD010440).

---

**Supplementary Figure 1:**
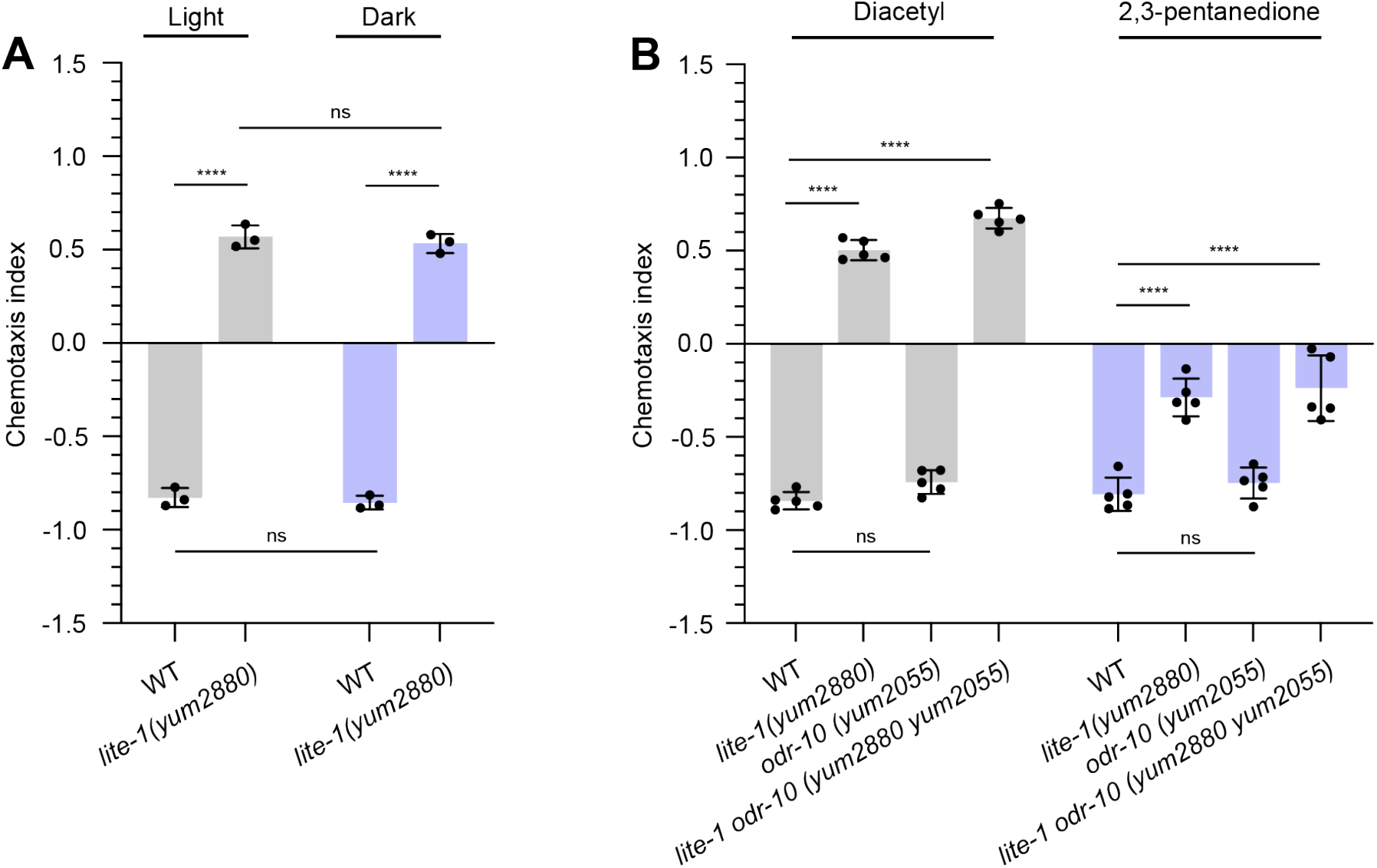
(A) Chemotaxis index of wild-type and *lite-1 (yum2880)* animals with undiluted diacetyl in the presence and absence of ambience light. P-values are from unpaired two-tailed t-tests, n=3 biological replicates, each an average of 3-4 plates. (B) Chemotaxis index of wild-type, *lite-1 (yum2880), odr-10 (yum2055) and lite-1odr-10 (yum2880; yum2055)* with undiluted diacetyl or 2,3-pentannedione. P-values are from unpaired two-tailed t-tests, n=5 biological replicates, each an average of 3-4 plates.

**Supplementary Figure 2:**
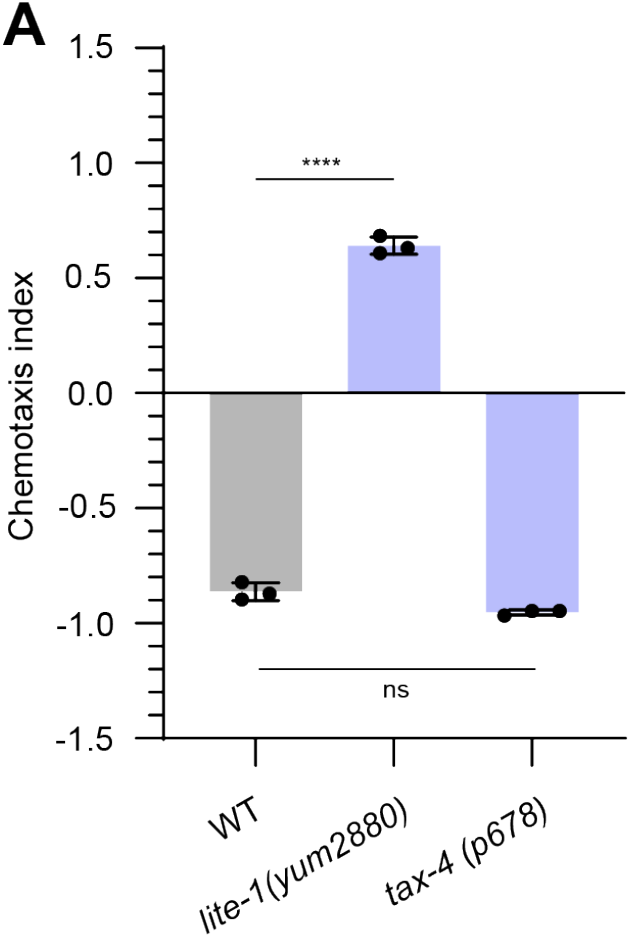
Chemotaxis index of wild-type and *tax-4 (p678)* animals with undiluted diacetyl. P-values are from unpaired two-tailed t-tests, n=3 biological replicates, each an average of 4 plates.

**Supplementary Figure 3:**
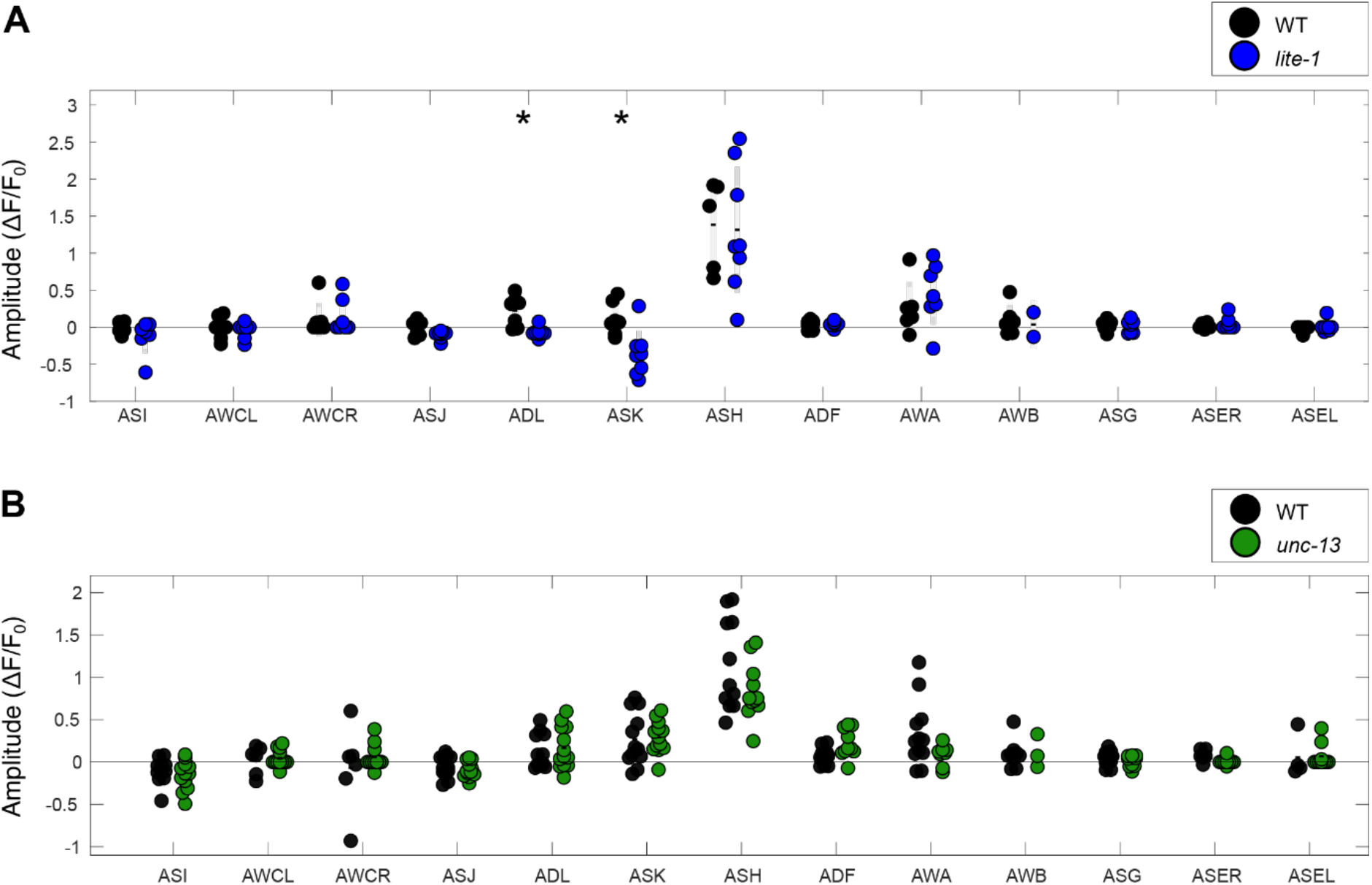
Scatter plot of calcium imaging responses across 11 pairs of sensory neurons following a pulse of diacetyl. (A) ADL and ASK neurons show significant differences in activity in the presence versus absence of LITE-1. (B) No significant differences in neuronal activity between wild-type and *unc-13* mutants across the different sensory neuron pairs. P-values were calculated using two-sided t-test with FDR correction, *p < 0.01.

**Supplementary Figure 4:**
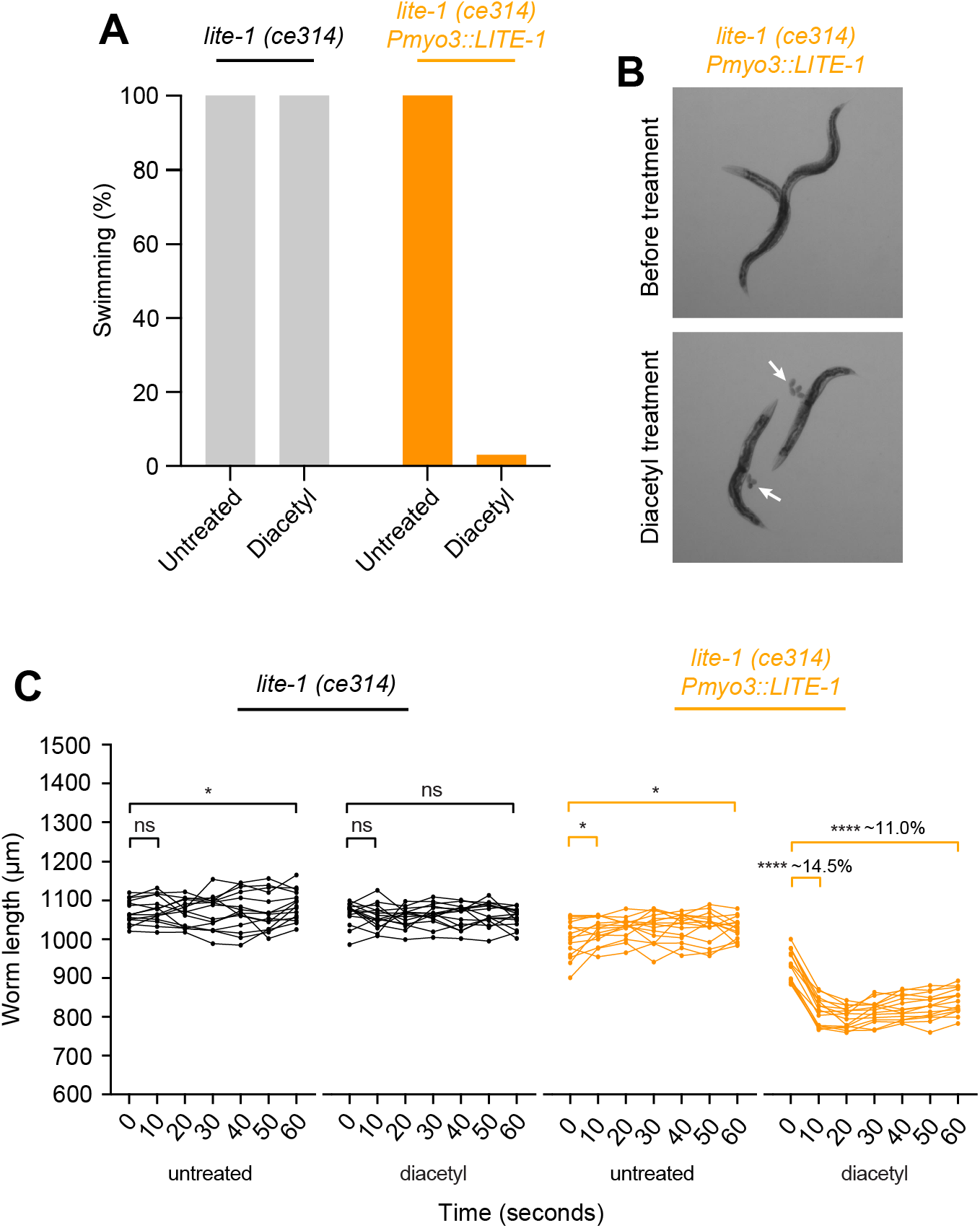
(A) Percent of worms that are swimming at ten seconds after exposure to buffer containing diacetyl (*lite-1 (ce314)*, left) whereas worms expressing LITE-1 in body-wall muscles (*lite-1 (ce314) Pmyo-3::*LITE-1, right) are mostly paralysed when treated with diacetyl. (B) Image of *lite-1 (ce314) Pmyo-3::*LITE-1 animals that ejected eggs after ten seconds of diacetyl treatment (white arrows). (C) Worm body length over time after treatment. *lite-1 (ce314)* worms expressing LITE-1 in body wall muscles show a significant contraction in response to diacetyl (right; insert showing the average contraction). P-values from paired two-tailed t-test, n ≥ 15 worms.

**Supplementary Figure 5:**
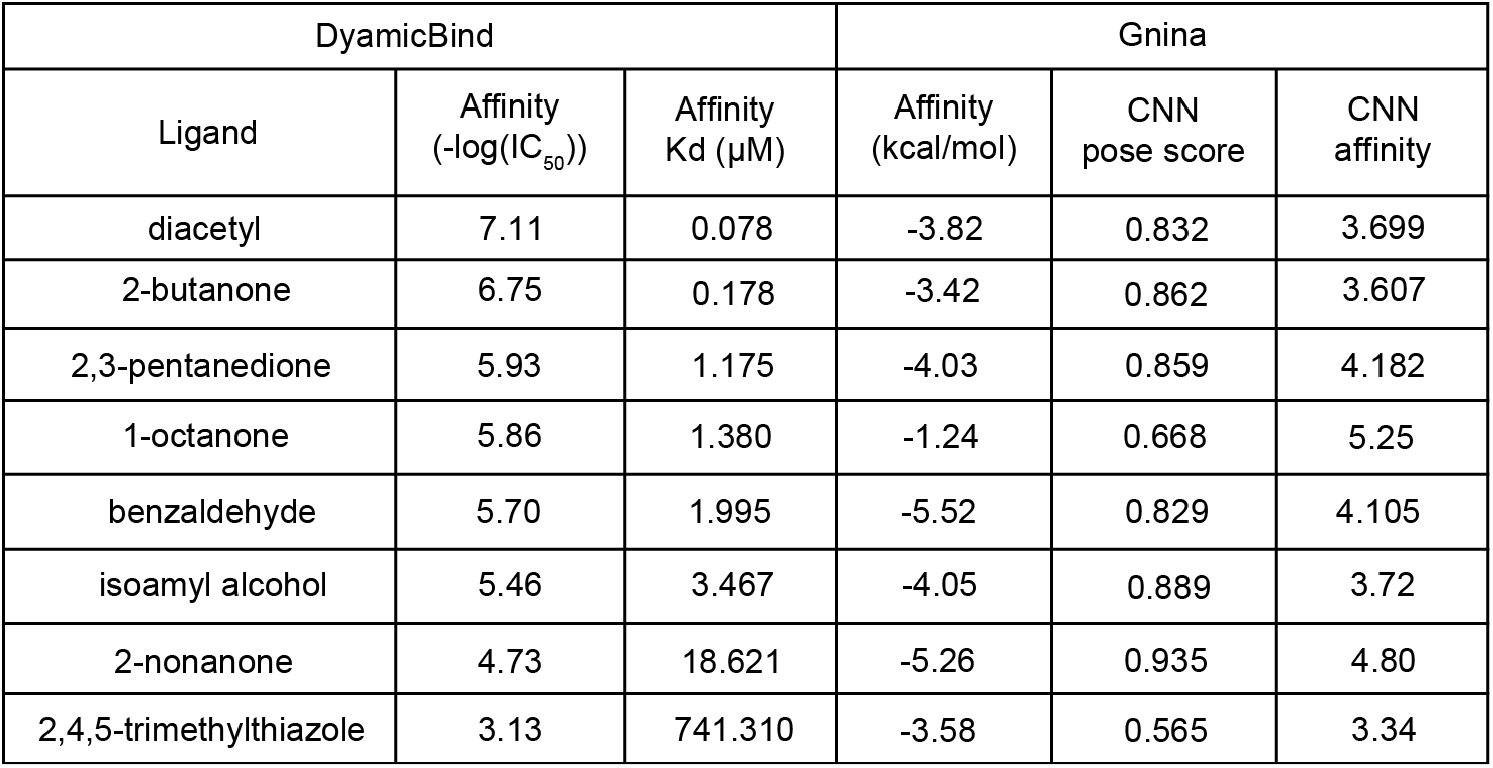
DynamicBind and Gninna predicted ligand binding affinities of different odorants to the putative LITE-1 binding pocket. For Dynamicbind, affinity is reported as -log(IC_50_), where higher values indicate stronger predicted binding and the corresponding Kd values(µM) were converted from - log(IC_50_), with lower values indicating stronger binding. For Gnina, affinity is reported as kcal/mol where higher values indicate higher binding interactions.

**Supplementary Figure 6:**
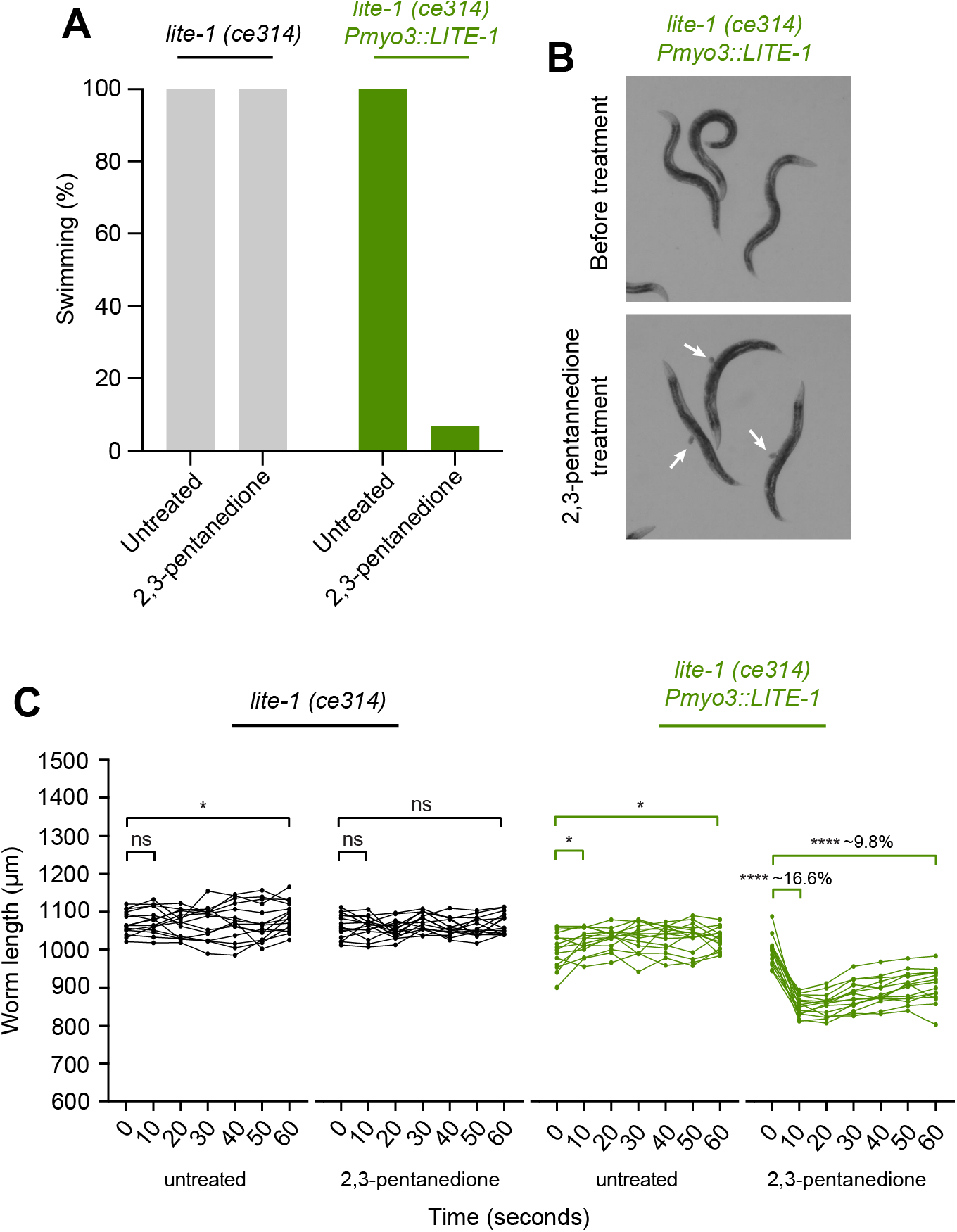
(A) Percent of worms that are swimming at ten seconds after exposure to buffer containing 2,3-pentanedione (*lite-1 (ce314)*, left) whereas worms expressing LITE-1 in body-wall muscles (*lite-1 (ce314) Pmyo-3::*LITE-1, right) are mostly paralysed when treated with 2,3-pentanedione. (B) Image of *lite-1 (ce314) Pmyo-3::*LITE-1 animals that ejected eggs after ten seconds of 2,3-pentanedione treatment (white arrows). (C) Worm body length over time after treatment. *lite-1 (ce314)* worms expressing LITE-1 in body-wall muscles show a significant contraction in response to 2,3-pentanedione (right; insert showing the average contraction). P-values from paired two-tailed t-test, n ≥ 15 worms.

**Supplementary video 1:** Chemotaxis assay without sodium azide showing avoidance of high concentration off diacetyl by wild-type animals. Wild-type animals were recorded over 1 h following addition of diacetyl and control to the chemotaxis plate. + indicates the position where 5 μl of undiluted diacetyl was spotted, which – indicates the position where 5 μl of ethanol was spotted (https://doi.org/10.6084/m9.figshare.32745054).

**Supplementary video 2:** Chemotaxis assay without sodium azide showing avoidance of high concentration off diacetyl by *lite-1* mutant animals. *lite-1* mutant animals were recorded over 1 h following addition of diacetyl and control to the chemotaxis plate. + indicates the position where 5 μl of undiluted diacetyl was spotted, which – indicates the position where 5 μl of ethanol was spotted (https://doi.org/10.6084/m9.figshare.32745054).

## References

Bargmann. (2006). Chemosensation in C. elegans

Bargmann, C. I., Hartwieg, E., & Horvitz, H. R. (1993). Odorant-selective genes and neurons mediate olfaction in C. elegans. Cell, 74(3), 515–527. 10.1016/0092-8674(93)80053-h

Bargmann, C. I., & Horvitz, H. R. (1991). Chemosensory neurons with overlapping functions direct chemotaxis to multiple chemicals in C. elegans. Neuron, 7(5), 729–742. 10.1016/0896-6273(91)90276-6

Barlow, I. (2019). Bleach Synchronisation of C. elegans v1. https://doi.org/10.17504/protocols.io.2bzgap6

Barlow, I. L., Feriani, L., Minga, E., McDermott-Rouse, A., O’Brien, T. J., Liu, Z., Hofbauer, M., Stowers, J. R., Andersen, E. C., Ding, S. S., & Brown, A. E. X. (2022). Megapixel camera arrays enable high-resolution animal tracking in multiwell plates. Commun Biol, 5(1), 253. 10.1038/s42003-022-03206-1

Bhatla, N., & Horvitz, H. R. (2015). Light and Hydrogen Peroxide Inhibit. Feeding through Gustatory Receptor Orthologs and Pharyngeal Neurons. Neuron, 85(4), 804–818. 10.1016/j.neuron.2014.12.061

Bischer, A. P., Baran, T. M., & Wojtovich, A. P. (2023). Reactive oxygen species drive foraging decisions in Caenorhabditis elegans. Redox Biol, 67, 102934. 10.1016/j.redox.2023.102934

Bokman, E., Kalij, I. P., & Zaslaver, A. (2024). Aberrant Positions of the Chemosensory Neurons in the Neurotransmitter-Release Mutant unc-13. Int J Mol Sci, 25(23). 10.3390/ijms252312956

Bokman, E., Pritz, C. O., Ruach, R., Itskovits, E., Sharvit, H., & Zaslaver, A. (2024). Intricate response dynamics enhances stimulus discrimination in the resource-limited C. elegans chemosensory system. BMC Biol, 22(1), 173. 10.1186/s12915-024-01977-z

Brenner, S. (1974). Genetics of Caenorhabditis-Elegans. Genetics, 77(1), 71–94.

Chronis, N., Zimmer, M., & Bargmann, C. I. (2007). Microfluidics for in vivo imaging of neuronal and behavioral activity in Caenorhabditis elegans. Nature Methods, 4(9), 727–731. 10.1038/nmeth1075

Cook, S. J., Jarrell, T. A., Brittin, C. A., Wang, Y., Bloniarz, A. E., Yakovlev, M. A., Nguyen, K. C. Q., Tang, L. T., Bayer, E. A., Duerr, J. S., Bulow, H. E., Hobert, O., Hall, D. H., & Emmons, S. W. (2019). Whole-animal connectomes of both Caenorhabditis elegans sexes. Nature, 571(7763), 63–71. 10.1038/s41586-019-1352-7

Dunkel, E., Aoki, I., Bergs, A., & Gottschalk, A. (2025). Neurons and molecules involved in noxious light sensation in Caenorhabditis elegans. G3 (Bethesda). 10.1093/g3journal/jkaf086

Edwards, S. L., Charlie, N. K., Milfort, M. C., Brown, B. S., Gravlin, C. N., Knecht, J. E., & Miller, K. G. (2008). A novel molecular solution for ultraviolet light detection in Caenorhabditis elegans. Plos Biology, 6(8), e198. 10.1371/journal.pbio.0060198

Gong, J. K., Yuan, Y. Y., Ward, A., Kang, L. J., Zhang, B., Wu, Z. P., Peng, J. M., Feng, Z. Y., Liu, J. F., & Xu, X. Z. S. (2016). The Taste Receptor Homolog LITE-1 Is a Photoreceptor. Cell, 167(5), 1252–+. 10.1016/j.cell.2016.10.053

Hanson, S. M., Scholuke, J., Liewald, J., Sharma, R., Ruse, C., Engel, M., Schuler, C., Klaus, A., Arghittu, S., Baumbach, F., Seidenthal, M., Dill, H., Hummer, G., & Gottschalk, A. (2023). Structure-function analysis suggests that the photoreceptor LITE-1 is a light-activated ion channel. Current Biology, 33(16), 3423–3435 e3425. 10.1016/j.cub.2023.07.008

Hanson, S. M., Scholüke, J., Liewald, J., Sharma, R., Ruse, C., Engel, M., Schüler, C., Klaus, A., Arghittu, S., Baumbach, F., Seidenthal, M., Dill, H., Hummer, G., & Gottschalk, A. (2023). Structure-function analysis suggests that the photoreceptor LITE-1 is a light-activated ion channel. Current Biology, 33(16). 10.1016/j.cub.2023.07.008

Hao, J. C., Adler, C. E., Mebane, L., Gertler, F. B., Bargmann, C. I., & Tessier-Lavigne, M. (2010). The tripartite motif protein MADD-2 functions with the receptor UNC-40 (DCC) in Netrin-mediated axon attraction and branching. Dev Cell, 18(6), 950–960. 10.1016/j.devcel.2010.02.019

Huang, C. Y., Li, J. B., Liu, W. B., & Li, C. J. (2019). Diacetyl as a “traceless” visible light photosensitizer in metal-free cross-dehydrogenative coupling reactions. Chemical Science, 10(19), 5018–5024. 10.1039/c8sc05631e

Itskovits, E., Ruach, R., Kazakov, A., & Zaslaver, A. (2018). Concerted pulsatile and graded neural dynamics enables efficient chemotaxis in C. elegans. Nature Communications, 9(1), 2866. 10.1038/s41467-018-05151-2

Iwanir, S., Ruach, R., Itskovits, E., Pritz, C. O., Bokman, E., & Zaslaver, A. (2019). Irrational behavior in C. elegans arises from asymmetric modulatory effects within single sensory neurons. Nature Communications, 10(1), 3202. 10.1038/s41467-019-11163-3

Javer, A., Currie, M., Lee, C. W., Hokanson, J., Li, K., Martineau, C. N., Yemini, E., Grundy, L. J., Li, C., Ch’ng, Q., Schafer, W. R., Nollen, E. A. A., Kerr, R., & Brown, A. E. X. (2018). An open-source platform for analyzing and sharing worm-behavior data. Nature Methods, 15(9), 645–646. 10.1038/s41592-018-0112-1

Lian, P. C., Li, R. Y., Wan, X., Xiang, Z. X., Liu, H., Cao, Z. Y., & Wan, X. B. (2022). Acetylation of alcohols and amines under visible light irradiation: diacetyl as an acylation reagent and photosensitizer. Organic Chemistry Frontiers, 9(2), 311–319. 10.1039/d1qo01613j

Liu, J., Ward, A., Gao, J., Dong, Y., Nishio, N., Inada, H., Kang, L., Yu, Y., Ma, D., Xu, T., Mori, I., Xie, Z., & Xu, X. Z. (2010). C. elegans phototransduction requires a G protein-dependent cGMP pathway and a taste receptor homolog. Nat Neurosci, 13(6), 715–722. 10.1038/nn.2540

Lu, W., Zhang, J., Huang, W., Zhang, Z., Jia, X., Wang, Z., Shi, L., Li, C., Wolynes, P. G., & Zheng, S. (2024). DynamicBind: predicting ligand-specific protein-ligand complex structure with a deep equivariant generative model. Nature Communications, 15(1), 1071. 10.1038/s41467-024-45461-2

McNutt, A. T., Francoeur, P., Aggarwal, R., Masuda, T., Meli, R., Ragoza, M., Sunseri, J., & Koes, D. R. (2021). GNINA 1.0: molecular docking with deep learning. J Cheminform, 13(1), 43. 10.1186/s13321-021-00522-2

Montell, C. (2009). A taste of the Drosophila gustatory receptors. Curr Opin Neurobiol, 19(4), 345–353. 10.1016/j.conb.2009.07.001

Pritz, C., Itskovits, E., Bokman, E., Ruach, R., Gritsenko, V., Nelken, T., Menasherof, M., Azulay, A., & Zaslaver, A. (2023). Principles for coding associative memories in a compact neural network. Elife, 12. 10.7554/eLife.74434

Pu, L. J., Wang, J., Lu, Q. X., Nilsson, L., Philbrook, A., Pandey, A., Zhao, L. A., van Schendel, R., Koh, A., Peres, T. V., Hashi, W. H., Myint, S. L., Williams, C., Gilthorpe, J. D., Wai, S. N., Brown, A., Tijsterman, M., Sengupta, P., Henriksson, J., & Chen, C. C. (2023). Dissecting the genetic landscape of GPCR signaling through phenotypic profiling in. Nature Communications, 14(1). https://doi.org/ARTN8410 z10.1038/s41467-023-44177-z

Quintin, S., Aspert, T., Ye, T., & Charvin, G. (2022). Distinct mechanisms underlie H2O2 sensing in C. elegans head and tail. PLoS One, 17(9), e0274226. 10.1371/journal.pone.0274226

Richmond, J. E., Davis, W. S., & Jorgensen, E. M. (1999). UNC-13 is required for synaptic vesicle fusion in C. elegans. Nat Neurosci, 2(11), 959–964. 10.1038/14755

Schindelin, J., Arganda-Carreras, I., Frise, E., Kaynig, V., Longair, M., Pietzsch, T., Preibisch, S., Rueden, C., Saalfeld, S., Schmid, B., Tinevez, J. Y., White, D. J., Hartenstein, V., Eliceiri, K., Tomancak, P., & Cardona, A. (2012). Fiji: an open-source platform for biological-image analysis. Nature Methods, 9(7), 676–682. 10.1038/Nmeth.2019

Sengupta, P., Chou, J. H., & Bargmann, C. I. (1996). odr-10 encodes a seven transmembrane domain olfactory receptor required for responses to the odorant diacetyl. Cell, 84(6), 899–909. 10.1016/s0092-8674(00)81068-5

Taniguchi, G., Uozumi, T., Kiriyama, K., Kamizaki, T., & Hirotsu, T. (2014). Screening of odor-receptor pairs in Caenorhabditis elegans reveals different receptors for high and low odor concentrations. Sci Signal, 7(323), ra39. 10.1126/scisignal.2005136

Toyoshima, Y., Tokunaga, T., Hirose, O., Kanamori, M., Teramoto, T., Jang, M. S., Kuge, S., Ishihara, T., Yoshida, R., & Iino, Y. (2016). Accurate Automatic Detection of Densely Distributed Cell Nuclei in 3D Space. PLoS Comput Biol, 12(6), e1004970. 10.1371/journal.pcbi.1004970

Ward, A., Liu, J., Feng, Z., & Xu, X. Z. (2008). Light-sensitive neurons and channels mediate phototaxis in C. elegans. Nat Neurosci, 11(8), 916–922. 10.1038/nn.2155

Zhang, W., He, F., Ronan, E. A., Liu, H., Gong, J., Liu, J., & Xu, X. Z. S. (2020). Regulation of photosensation by hydrogen peroxide and antioxidants in C. elegans. PLoS Genet, 16(12), e1009257. 10.1371/journal.pgen.1009257

Zhang, X., Liu, J., Pan, T., Ward, A., Liu, J., & Xu, X. Z. S. (2022). A cilia-independent function of BBSome mediated by DLK-MAPK signaling in C. elegans photosensation. Dev Cell, 57(12), 1545–1557 e1544. 10.1016/j.devcel.2022.05.005

Zhang, Y., Chou, J. H., Bradley, J., Bargmann, C. I., & Zinn, K. (1997). The Caenorhabditis elegans seven-transmembrane protein ODR-10 functions as an odorant receptor in mammalian cells. Proc Natl Acad Sci U S A, 94(22), 12162–12167. 10.1073/pnas.94.22.12162

